# Filamentation-driven peripheral clustering of the inducible lysine decarboxylase is crucial for *E. coli* acid stress response

**DOI:** 10.1101/2025.03.27.645680

**Authors:** Moritz A. Kirchner, Jessica Y. El Khoury, Frédéric Barras, Jean-Philippe Kleman, Irina Gutsche

## Abstract

Bacteria have evolved numerous sophisticated acid stress response strategies to withstand fluctuating environmental pH, with enterobacterial inducible amino acid decarboxylases playing a major role. The lysine decarboxylase LdcI catalyses lysine-to-cadaverine conversion coupled to proton consumption and carbon dioxide release, thereby buffering cytoplasmic and extracellular pH. Our previous study showed that *Escherichia coli* LdcI forms intracellular patches under mild acid stress, and that at the corresponding acidic pH, purified LdcI polymerises into filaments, suggesting a potential role for polymerisation in its function. Here, we investigate the physiological relevance of LdcI filamentation in *E. coli* using 3D super-resolution fluorescence microscopy and a structure-based LdcI polymerisation-deficient *E. coli* mutant strain. We establish a semi-automated workflow for detection of clusters within a bacterial cell as well as quantitative analysis of their volume and localisation. We demonstrate that LdcI predominantly clusters at the cell periphery. We then show that disrupting LdcI polymerisation markedly reduces cluster size without significantly affecting localisation, suggesting that acid-stress induced LdcI clustering *in vivo* is driven by the enzyme’s filamentation. Growth and pH measurements reveal that the mutant strain exhibits reduced fitness and impaired extracellular buffering compared to the wild type, indicating that LdcI polymerisation enhances *E. coli* acid stress response. Our findings provide strong evidence that LdcI filamentation is a key regulatory feature of the *E. coli* capacity to counteract acid stress, optimising enzymatic activity by spatially organising LdcI within the cytoplasm. More broadly, this work adds to a growing body of evidence supporting the functional significance of metabolic enzyme self-assembly and its implications for bacterial stress adaptation.

**Author summary:** Amino acid decarboxylases confer enterobacteria the ability to survive under acidic conditions. Kirchner et al. show that polymerisation of the acid-stress inducible lysine decarboxylase LdcI within *E. coli* cells leads to its peripheral intracellular clustering and fine-tunes its enzymatic activity *in vivo*.

## Introduction

Pathogenic and commensal bacteria have evolved a diverse array of strategies to survive and even flourish in adverse environments. One common challenge they often face is acidity, encountered in external settings such as soil and food, as well as within the host’s dental plaque, gastrointestinal tract, and macrophage phagosomes [1]. To counteract this stress, many neutrophiles depend on inducible acid resistance systems, which employ proton-consuming amino acid decarboxylases and cognate antiporters [2, 3]. These systems import amino acid substrates - primarily glutamate, arginine or lysine - in exchange for more alkaline polyamine products, effectively buffering the bacterial cytosol and the local extracellular environment.Beyond acid stress response, polyamines contribute to a broad spectrum of processes including bacterial physiology, gene regulation, biofilm formation, virulence and antibiotic resistance [4]. *Escherichia coli* possess all three major inducible amino acid decarboxylase systems, predominantly conserved in *Enterobacteriaceae* [5]. Each system is finely tuned for a certain strength of acid stress and depends on environmental conditions and growth phase. In addition, their activation has recently shown to be distributed across individual cells in the population, creating a “division of labor” that ensures global fitness over a wide range of low pH conditions [6].

Under moderate acid stress in a lysine-rich environment, the majority of *E. coli* cells activate the lysine-to-cadaverine converting cad system, composed of the *cadBA* operon and its transcription regulator *cadC* [3, 7]. In non-inducing conditions, the membrane-integrated CadC is inhibited by binding to the lysine-specific membrane permease LysP [8]. Lysine import by LysP under acidic conditions releases CadC, enabling its activation through dimerisation of the periplasmic sensory domain and structural rearrangements in the cytoplasmic DNA-binding domain, along with reduction of a disulfide bond and subsequent proteolytic cleavage by periplasmic proteases (REFS). Activated CadC diffuses within the membrane and binds the *cadBA* promoter, inducing production of the lysine decarboxylase CadA and the inner membrane lysine/cadaverine aniporter CadB, until CadC inhibition by excess of the exported cadaverine shuts down the system [9, 10]. CadA, termed LdcI for acid stress-inducible lysine decarboxylase, is a dihedral symmetric enzyme extensively studied since the 1940s due to its role in acid stress response related to enterobacterial pathogenicity and the numerous clinical, agricultural, and industrial applications of bio-based cadaverine [3, 7, 11–13].

An increasing number of metabolic enzymes, particularly those with a dihedral symmetry, are found to self-assemble into filaments [14–16]. Exploring the functional and regulatory benefits of enzyme filamentation is becoming a prominent research area. Our previous cryo-EM analysis has revealed that at pH 5.7 - optimal for *E. coli* LdcI expression and activity - D5-symmetric LdcI stacks into filamentous polymers *in vitro* [17]. Corroborated by theoretical calculations of the charge distribution and the folding energy as a function of pH, the stacking propensity of LdcI at pH below 6 is very high, while no filamentation occurs above this threshold [18]. Furthermore, the structural determinants of the LdcI stacking are conserved across *Enterobacteriaceae*, suggesting that acid stress-induced LdcI polymerisation may be a common mechanism in this bacterial family [18].

In parallel to growing interest in mechanistic and structural investigation of metabolic enzyme polymerisation *in vitro*, advances in light microscopy [19], and since very recently, also cryo-electron tomography [20], prompt to correlate this phenomenon with punctate or filamentous assemblies *in vivo*. It becomes increasingly apparent that cells biochemically and spatially regulate numerous enzymes through stress-triggered condensation into liquid droplets, amyloid-like aggregates or ordered filaments [21]. Acidification of the growth medium is one such stressor, and indeed, our earlier immunofluorescence imaging showed a patchy distribution of LdcI in acid-stressed *E. coli* [17]. Since the pH that induces LdcI filamentation *in vitro* matches the internal pH in the *E. coli* cell upon acid stress, we have hypothesised that the acidification-induced LdcI patches *in vivo* represent LdcI stacks formed to locally increase the enzyme concentration and thereby boost its activity [17].

Although logically sound, this hypothesis requires experimental validation. To this end, we have engineered a structure-based triple triple mutant of *E. coli* LdcI that loses polymerisation capacity *in vitro* and remains decameric even at low pH. We have incorporated the modified cognate allele into the *E. coli* chromosome, generating an *E. coli* mutant, hereinafter termed 3M, in which the LdcI filamentation is expected to be compromised. We have then quantitatively compared the predominantly peripheral intracellular distribution and the acid stress-induced clustering of endogenous LdcI in the wild type MG1655 (WT) and triple mutant (3M) *E. coli* strains. The results provide compelling evidence that the *E. coli* 3M strain exhibits smaller LdcI clusters than those of the WT strains, confirming our initial intuition. Furthermore, the 3M mutant is less efficient in counteracting acid stress in terms of growth and external pH buffering, demonstrating that polymerisation is indeed required for optimal LdcI enzyme activity *in vivo*. Altogether, these findings lead us to conclude that LdcI clustering *in vivo* is driven by its filamentation, required for optimal acid stress response in *E. coli*.

## Materials and methods

### Construction of the chromosomal mutant *ldcI-3M* containing the following three mutations: E445A-D447A-R448E (FBE 116 strain)

Firstly, the gene ”ldcI E445A-D447A-R448A”, called `ldcI_3M`, was amplified by PCR using the plasmid “pET22b-LdcI-His E445A-D447A-R448A” as template. This amplicon was then cloned into the plasmid pJL72 KanR. Secondly, this new plasmid (pJEK02 (pJL72 + `ldcI_3M`)) was used to amplify the *ldcI-3M* gene plus the kanamycine resistance cassette. Thirdly, a transformation of this amplicon into the FBE832 (`MG` Δ`ldcI::FRT`) strain was performed using the Datsenko and Wanner technique [22]. Finally, the kanamycine resistance cassette was removed by transformation with pCp20. Strains, plasmids and primers used to prepare this mutant are listed in Suppl. File 1.

### Cell culture

A 350 ml liquid LB culture was inoculated with an overnight preculture of *E. coli* MG1655 (either WT or harbouring the 3M LdcI variant) to a final OD of 0.01 and grown at 37°C. At OD 0.1 cells were harvested by centrifugation (20°C, 4500 g, 20 min) and resuspended in the same volume of LB containing 30 mM L-lysine and pH-adjusted to pH 4.6 (HCl). The culture volume was then split into batches from which cells were harvested at 0, 30, 60, 90 and 180 minutes by centrifugation (4°C, 4000 g, 10 min) to give an amount which corresponds to an OD 1 resuspended in 1 ml for western blot analysis (see below). At 90 min, additional samples were harvested by centrifugation (OD 4 for 1 ml) for fluorescence imaging. Cell growth (OD) and pH of each batch were measured at the time points stated above for the harvest.

### Sample preparation for fluorescence microscopy

For imaging, samples were fixed using a 4 % formaldehyde solution in PBS by constant agitation at room temperature for 45 min. After washing with PBS by centrifugation (20°C, 1620 g, 10 min), cells were permeabilised using 0.1 % Triton X-100 in PBS for 10 min at room temperature, and then washed with PBS by centrifugation. The cell suspension was then incubated with 1 % BSA in PBS for 30 min at room temperature before 0.5 µg of camelid nanobody coupled to AlexaFluor™647 (Thermo Fisher Scientific, USA) was applied as previously described [17]. In brief, cell suspension was incubated with the nanobody solution for 16 h at 4°C. After incubation, the cells were washed thrice with PBS to remove unbound nanobody. Cover slips were cleaned by UV irradiation for 15 min on either side, before a 10 µl drop of cell suspension was placed in the centre of the cover slip and the suspension was left to dry. A 10 µL drop of STORM buffer containing GLOX solution [23] was placed on top. Finally a 1.5 % low-melting agarose pad was cast between cover slip and slide for cell immobilisation of cells during imaging.

### 3D Single Molecule Localisation Microscopy (SMLM)

SMLM was performed using direct stochastic optical reconstruction microscopy (dSTORM) [24, 25] using the Abbelight™SAFe360 setup of the cell-imaging platform (M4D) composed of an inverted microscope (Olympus IX83, Olympus, Japan) in combination with an L6Cc laser combiner (Oxxius, France). Excitation laser light was focused onto the back focal plane of the objective lens (Oil Immersion 100×, NA 1.5). All streams were recorded in HILO illumination mode [26]. Samples were constantly illuminated with laser light at 642 nm over a region of 70x70 µm at a power density of 637 W/cm^2^. For fluorophore reactivation, we used gradually increasing power of a 405 nm laser beam to illuminate the sample region. Stream acquisitions were done using Hamamatsu Orca Fusion sCMOS cameras for 40,000 frames at 27°C and an exposure time of 50 ms. To achieve three-dimensional signal detection, a cylindrical lens was introduced into the detection light path [27]. PSF fitting was performed using the Abbelight™NEO analysis software using a maximum likelihood protocol and cross-correlation based drift correction after any 1,000 frames [28]. Localisations with a higher axial uncertainty than 25 nm were removed from the final table.

### Clustering analysis

Clustering analysis was performed on single cells segmented from the full recorded volume. In order to segment the single cells, the DBSCAN algorithm was used as implemented in Abbelight™NEO analysis [29]. For cells which appeared to have recently divided, accurate segmentation by clustering was not always possible. In these cases, the resulting two-cell files were only used for cluster volume determination and not for distance ratio calculations (see below). Single cell data were tested for significant clustering behaviour using analysis with Ripley’s function as implemented in locan [30, 31]. The single cell files were used for clustering using FOCAL3D [32]. Clustering parameters were individually determined for each single cell. Coordinates for individual clusters were extracted and the volume calculated by determining the volume of a convex hull around all points by defining the outer most points as vertices for the hull. Coordinates of individual clusters were used to determine the position of centroids of clusters relative to the cell centre (centre of the cloud of all points). The set of points defining the boundary of the point cloud was found by building a convex hull around all points. Distances between cluster centroids and the cell centre *d*_*c*_, as well as distances between cluster centroids and the closest boundary point of the entire point cloud *d*_*v*_ were determined. The distance ratio *r*_*c*_ of a centroid was defined as:

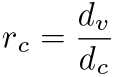

The distances were also determined for *n* randomly created test centroids following a uniform distribution in the cell volume, where *n* is the number of points found in the point cloud of the cell. The distance of all test centroids to the cell centre *d*_*tc*_ and to the closest boundary point *d*_*tv*_ were determined. Distances of all test centroids were averaged to give the average distance ratio 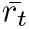. A cluster was defined as peripheral if this average ratio was found to be greater than the individual ratio for a single real cluster centroid:

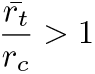

### Western-blotting

Cells were harvested by centrifugation as described above and resuspended in WB-buffer containing 1X Laemmli buffer (Bio-Rad, USA) and 2.5 % beta-mercaptoethanol in 1X TBS (Euromedex, France) Samples were run on a 12 % acrylamide SDS-PAGE gel before western-blotting on nitrocellulose (semi-dry blotting following manufacturer instructions). Nitrocellulose membrane was blocked with 5 % BSA in TBS-T (0.1 % Tween20 in 1X TBS) overnight at 4°C. After washing with TBS-T the membrane was incubated with anti-Ldc antibody, washed again and then incubated with a secondary anti-rabbit antibody (Sigma-Aldrich, USA). Signal was detected on the membrane after treatment with WB-substrate (Lumi-Light Western Blotting Substrate, Roche, Switzerland).

## Results

### Heterogeneous expression of LdcI in acid-stressed *E. coli* complicates 3D single molecule imaging

In our previous work we introduced and validated a procedure of immunofluorescence LdcI imaging in acid-stressed *E. coli* with a help of an anti-LdcI nanobody, labelled with the AlexaFluor™647 dye (AF647) [17]. To our surprise, endogenous LdcI, produced 90 minutes after acidification of the lysine-containing LB growth medium to pH 4.6 in oxygen-limiting conditions to counteract acid stress, appeared to form patches, which we then hypothesised to correspond to LdcI stacks. To obtain quantitative insights into the LdcI localisation and its potential relation to the propensity of LdcI to polymerise at low pH, we now set out to characterise the intracellular distribution of LdcI using three-dimensional stochastic optical reconstruction microscopy (3D-STORM) and collected extensive data to enable not only single cell but also population-level quantification and conclusions.

As previously reported and contrary to what might be expected from a highly soluble protein, LdcI generally did not show a uniform distribution throughout the cytoplasm of a single cell but formed variably-sized higher-density patches (Fig 1A, Suppl. Fig. 1). However, quantitative analysis of these patches was not straightforward for the following reasons. First, the average signal density varied greatly between cells, which is most plausibly explained by the recently discovered heterogenous expression of the transcription regulator CadC across the *E. coli* population [33], although a possibility of an incomplete permeabilisation cannot be fully excluded. This led in particular to a considerable number of ”dark” cells with a very low overall signal density, which were almost invisible in the immunofluorescence images (Fig 1A) and therefore had to be excluded from further processing for the sake of accuracy. Second, in the “bright” cells, the signal density was often very strong throughout the whole cytoplasm, certainly due to a massive induction of the LdcI production in response to acid stress. Such high signal posed a substantial complication in detecting regions of concentrated signal, which reflect elevated protein density. Thus, to enable rigorous characterisation of the LdcI distribution, we proceeded with the development of an efficient and unbiased workflow for semi-automatic detection of LdcI clusters and quantitative analysis of their volume and cellular localisation.

**Fig 1.**
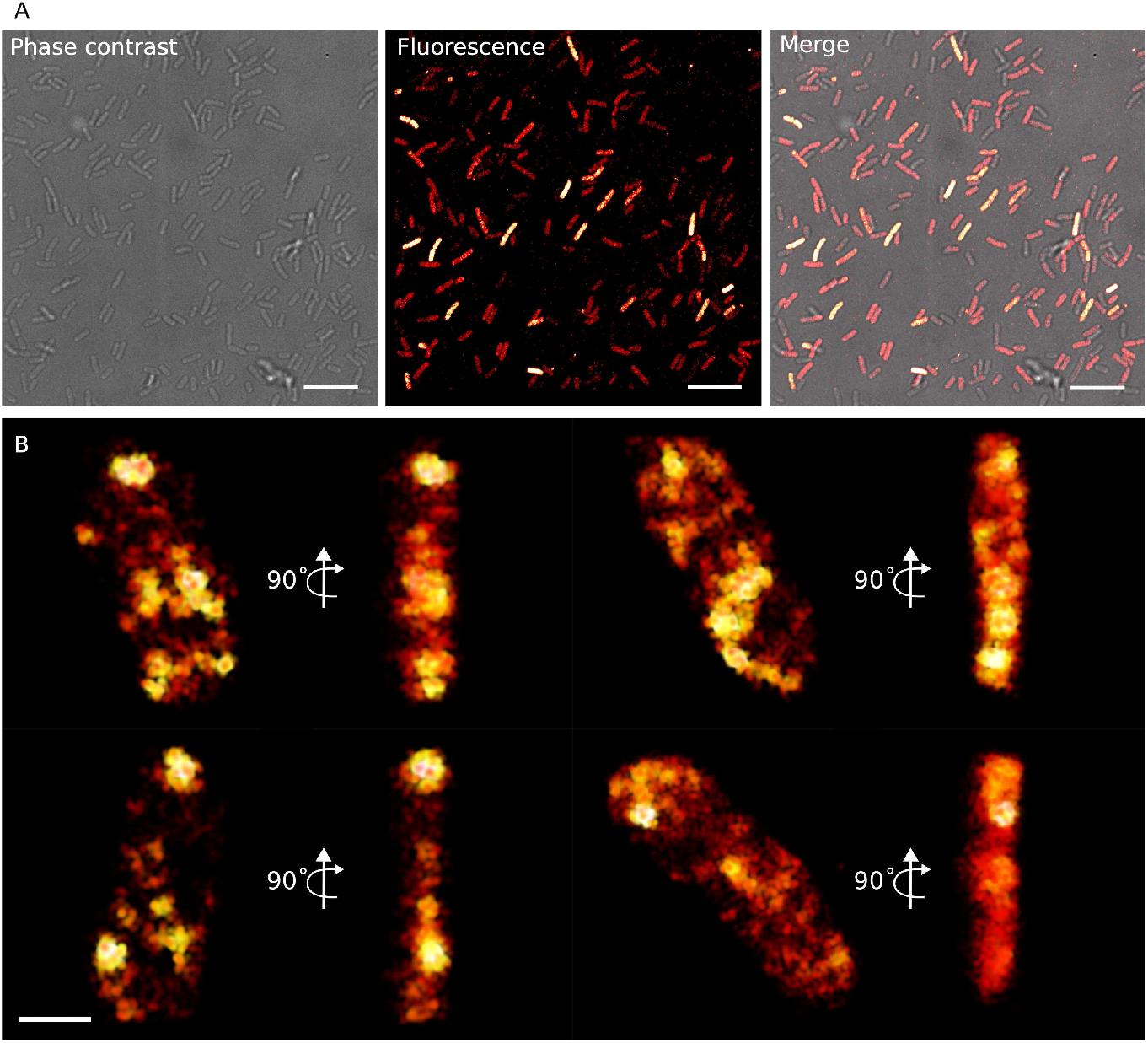
Three dimensional dSTORM imaging of LdcI in acid-stressed *E. coli*. Streams were recorded 90 min after exposure to pH 4.6 to induce LdcI expression. LdcI was visualised by immunofluorescence labelling with a camelid anti-LdcI nanobody coupled with AlexaFluor™647. A: A representative phase contrast image, a fluorescence image reconstructed using the acquired streams acquired and their merge are shown from left to right, to illustrate the differential expression of LdcI across the bacterial population. Scale bar = 10 µm. B: Four single cells (from multiple acquisitions) segmented from the full reconstructed 3D volume are shown in two different orientations. Brightness of points is proportional to the local signal density. Scale bar = 1 µm.

### Quantification of intracellular LdcI localisation in WT *E. coli* reveals predominant peripheral clustering without long-range arrangement or preference toward cell poles

The determination of the exact localisation and dimensions of the patches observed in the 3D dSTORM data was first explored using multiple standard clustering algorithms such as Voronoi tessellation [34, 35], multi-distance spatial cluster analysis based on Ripley’s K-function [30], and density-based clustering using DBSCAN (Density-Based Spatial Clustering of Applications with Noise) [29] (for a comprehensive review of clustering methods in single molecule localisation microscopy see [36]). After comparison, we opted for density-based methods. While the DBSCAN algorithm adequately segmented single cells from the full recorded volume, it was insufficient for accurately detecting discrete clusters within individual cells. Instead, intracellular clustering was successfully achieved with FOCAL3D [32], a DBSCAN-based 3D algorithm optimised for single-molecule localisation microscopy (SMLM), which builds clusters from subvolumes populated with data points rather than relying on radial distances between individual points. Using FOCAL3D, we successfully detected well-separated clusters of LdcI within the cells. Additionally, we expanded the data processing pipeline to calculate the volume of individual clusters by determining the volume inside a convex hull constructed from the outermost points of each cluster. Furthermore, we implemented a step to quantify whether a cluster localises to the cell periphery (see Methods) by comparing between the relative position of its centroid to the cell periphery and centre with the average of *n* = (number of data points) test centroids generated within the cell volume according to a uniform distribution. The full workflow is illustrated in Fig 2A. Clustering analysis was performed using only data points from single cells segmented from the full recorded volume.

**Fig 2.**
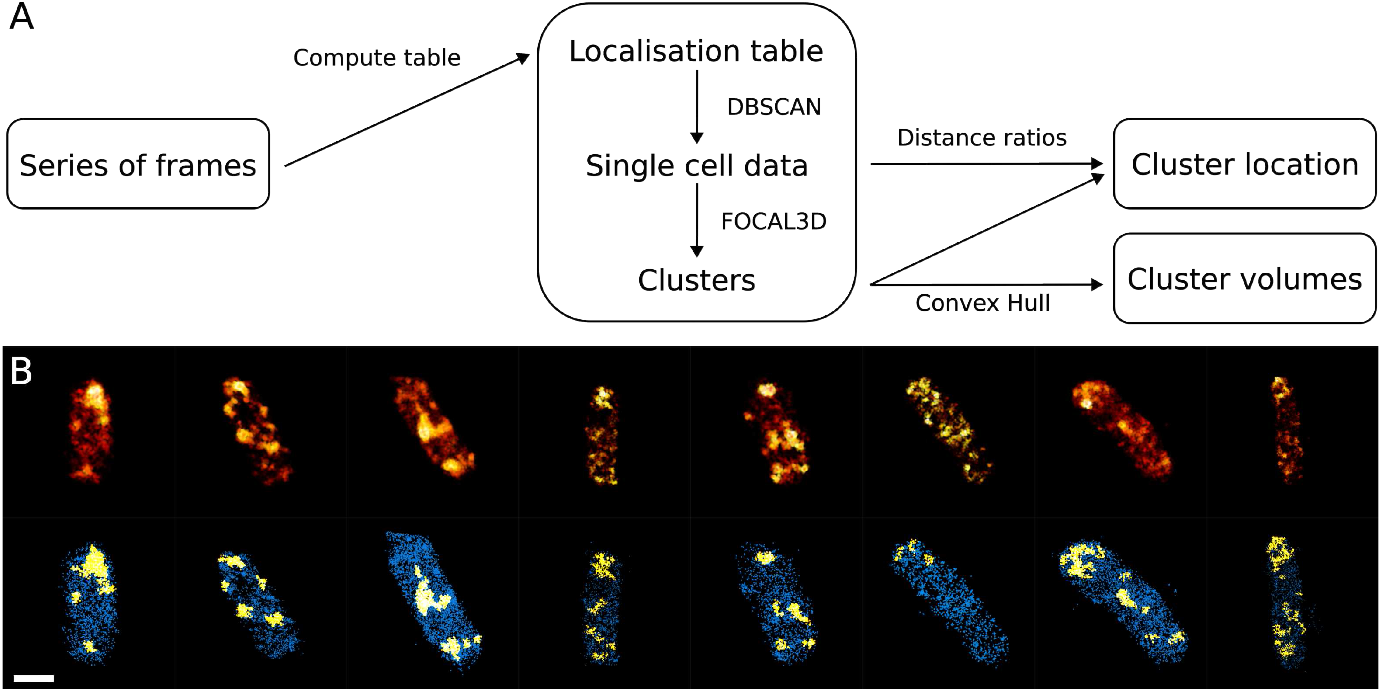
Workflow used for LdcI cluster identification and analysis. A: Flow chart of the clustering workflow. Compute table describes PSF fitting and drift correction of the streams recorded to compute the table storing all point coordinates and statistics. See Methods for explanation on calculation of distance ratios. B: Gallery of WT MG1655 *E. coli* cells segmented from several volumes using DBSCAN, which have been subjected to clustering analysis with FOCAL3D. The lower row shows the same cells as the upper row, with all localisations sorted into clustered (yellow) and non-clustered (blue). Scale bar = 1 µm.

Application of the developed pipeline to acid-stressed WT *E. coli* cells revealed the LdcI distribution within these single cells (Fig 2B). 353 individual clusters were extracted from 71 cells, with 226 clusters showing a centroid position closer to the cell periphery than the average test cluster centroid, accounting for 64 % of all observed clusters. Thus, we conclude that although, after its acid stress-induced production, LdcI can be found throughout the bulk of the *E. coli* cytoplasm, the enzyme preferentially clusters at the cell periphery, most likely underneath the inner membrane. Interestingly, no well-separated subpopulation of inter-point distances between cluster centroids per cell could be identified (Suppl. Fig. 2), indicating that the distribution of the LdcI clusters shows no considerable bias towards the cell poles. In addition, the resulting cluster distributions do not corroborate our initial visual impression - based on a limited 3D dSTORM dataset and lacking the present quantitative assessment - of a potential long-range pattern in cluster organisation.

### Disrupting LdcI polymerisation impedes *E. coli* acid stress response

Having set up the cluster analysis pipeline for 3D dSTORM imaging of endogenous acid stress-induced LdcI in the WT *E. coli* cells, we were ready to investigate the effects of disruption of the LdcI filamentation on the LdcI distribution. Thanks to our previous structural characterisation of the *E. coli* LdcI polymers and the phylogenetic analysis of bacterial Ldcs, we identified the conserved LdcI residues involved in linear stacking of *Enterobacteriacea* LdcI decamers [17]. Since the triple E445A-D447A-R468A mutation of *E. coli* LdcI abolishes decamer stacking *in vitro*, we engineered a mutant 3M strain harboring these three mutations in the *cadA* gene. A pH shift experiment described in our previous work (see also Methods) was performed on the WT and the 3M strains in parallel to evaluate potential effects of the *ldcI* mutations on the bacterial growth after acid stress, the concominant LdcI production and the resulting buffering of the extracellular medium. As shown in Suppl. Fig. 3A, while the growth curves of the WT and the 3M cells did not significantly differ during the first hour and a half after initiation of acid stress by transfer of the bacteria from pH 7.0 to 4.6, the 3M cells progressively lagged behind the WT and did not reach the same cell density as the WT cells after three hours of growth. Moreover, the pH of the culture medium remained systematically lower, showing that the buffering capacity of the 3M strain was considerably less efficient than that of the WT strain Suppl. Fig. 3B. After three hours post pH shift, in contrast to the WT strain that returned to pH 6.9, the pH of the 3M strain growth medium was still only around 6.2, although the steady state level of LdcI as assessed by western blot detection in full cell extracts was not affected by acid stress (Suppl. Fig. 3C,D). Consequently, we conclude that disrupting LdcI polymerisation markedly compromises the ability of *E. coli* to withstand acid stress. The negative effect on the phenotype was already evident after 90 min of growth, the time point at which the cells were harvested for immunofluorescence imaging.

### Disrupting LdcI filamentation reduces the size of intracellular LdcI clusters

The 3M strain was prepared and imaged similarly to the WT strain and the 3D dSTORM data processed using the above-described pipeline. As for the WT, the cells manifested high signal variability (Fig 1A, Suppl. Fig. 4), with “dark” cells being excluded from further analysis, and “bright” cells often showing high signal density throughout the volume (Fig 3, Suppl. Fig. 1). Although quantitative localisation analysis confirmed the initial visual impression of the absence of very large patches in the 3M cells (Fig 3B), the data processing pipeline showed that the polymerisation-preventing LdcI mutations did not completely abolish the enzyme clustering. Remarkably however, the average volume of the LdcI clusters was about 2.6 times smaller in the mutant bacteria as compared to the WT (Fig 3A). These smaller clusters are still preferentially located at the cell periphery - of 248 clusters observed in 65 individual cells, 167 had centroids at locations closer to the periphery than the average test centroid, corresponding to 68 % of all clusters. Thus, hindering acid-driven LdcI filamentation affects bacterial performance in the acid stress response and LdcI cluster size, without notably altering their overall distribution.

**Fig 3.**
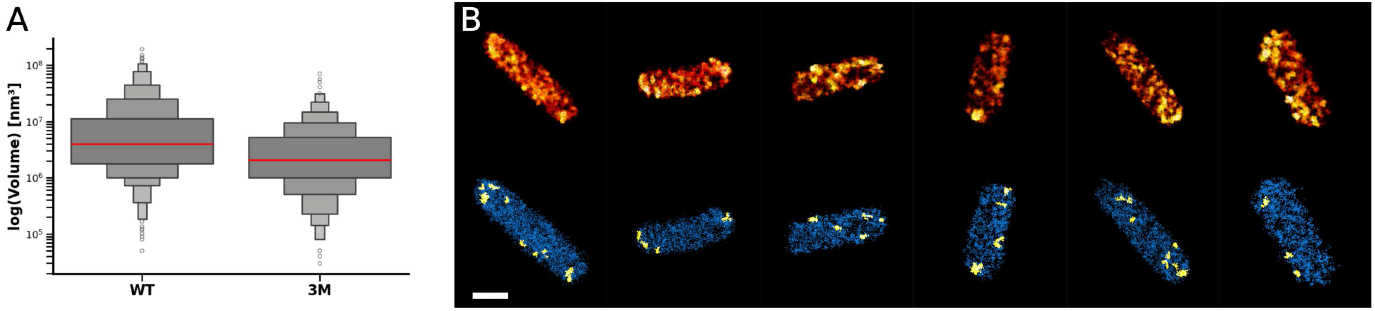
Residual LdcI clustering observed in the *E. coli* triple mutant strain. A: Boxenplot showing the difference in volume distribution between clusters detected in *E. coli* WT and 3M cells. B: Gallery of 3M cells segmented from several volumes using DBSCAN, which have been subjected to clustering analysis with FOCAL3D. The lower row shows the same cells as the upper row, with all localisations sorted into clustered (yellow) and non-clustered (blue). Scale bar = 1 µm.

## Discussion

While the past decade has brought significant advances in understanding the enterobacterial LdcI-based acid stress response, the field still resembles a puzzle, with pieces tentatively assembled but lacking some critical fragments needed to form a cohesive whole. Multiple structures of the decameric D5-symmetric *E. coli* LdcI and its homologues across diverse bacterial families captured these proteins in various functional states [17, 37–39]. The pH-dependent activity of the *E. coli* enzyme was determined to be optimal at pH 5.7, mirroring the internal pH of the bacterium under mild acid stress that induces *ldcI* gene expression [12]. At this same pH, LdcI decamers were shown to polymerise into filamentous stacks *in vitro* [12, 17], joining a growing number of dihedral symmetric enzymes exhibiting similar behavior. Indeed, dihedral symmetry is recognised to provide an optimal framework for the evolution of filamentation through a minimal number of mutations, enabling dynamic and reversible regulation of protein function via modulating structural stability, activity and spatial localisation in response to environmental cues [40, 41]. LdcI is a compelling example of these principles, with the molecular determinants of its polymerisation and the key pH-sensitive histidine residue conserved across enterobacteria [18], underscoring the evolutionary and functional significance of LdcI filaments. Therefore, as our initial immunofluorescence imaging indicated that LdcI formed peripheral patches in acid-stressed *E. coli*, it was reasonable to suggest that these patches corresponded to intracellular clusters of LdcI polymers crucial for optimal enzymatic activity and conferring enterobacteria a functional advantage in acid stress response [17]. However, testing this hypothesis required addressing three major gaps. First, 3D SMLM imaging of a large number of bacteria coupled with rigorous quantitative analysis was essential to obtain reliable information on LdcI distribution within individual cells and across the bacterial population. Second, a mutant *E. coli* strain retaining WT regulation of LdcI expression but with impaired LdcI polymerisation had to be engineered, to assess the importance of polymerisation for counteracting acid stress. Third, it was necessary to investigate whether polymerisation-deficient LdcI could still form intracellular clusters similar to those of the WT enzyme, to elucidate the anticipated relationships between filament assembly and cellular localisation.

Our comparative analysis of the growth and the external pH curves between the WT *E. coli* MG1655 strain and its chromosomal variant harboring a mutated LdcI version designed to keep the enzyme in its decameric state, revealed reduced fitness and buffering capacity of the LdcI variant synthesising 3M strain in a pH shift experiment. These results, combined with earlier data [17], demonstrate that LdcI polymerisation enhances the enzyme’s activity within the cell as a direct response to the environmental pH drop, helping the bacterium to efficiently counteract the acid stress. Curiously, the 3M LdcI strain still exhibited intracellular clustering despite its impaired polymerisation capacity, observed *in vitro* at low protein concentrations. This suggests the mutant decamer may retain residual stacking ability *in vivo*, in regions of particularly high local concentration. Importantly, quantitative clustering analysis revealed significantly smaller cluster volumes in the mutant compared to the WT strain (Fig 3). This strongly supports the hypothesis that intracellular LdcI clusters observed in acid-stressed *E. coli* cells are primarily composed of LdcI filaments. Together, our past and current work provides convincing evidence that *E. coli* response to mild acid stress that leads to the intracellular pH optimal for enzymatic activity of LdcI, triggers its expression and polymerisation, resulting in its intracellular clustering.

While the molecular mechanisms and functional consequences of enzyme polymerization are increasingly studied *in vitro* using cryo-EM combined with biochemical and biophysical analyses, the effects of filamentation-affecting mutations on endogenous enzyme function and distribution *in vivo* remain underexplored, particularly in bacteria. This is partly due to the scarcity of optical imaging studies on bacterial enzymes, whether they form filaments or not, as the small size of bacteria necessitates SMLM to achieve sufficient resolution [19]. Additionally, the two fields - *in vitro* mechanistic studies of enzyme filamentation and *in vivo* investigations of its functional consequences - rarely intersect. Our work highlights how identification of the structural underpinnings of LdcI polymerisation leads to crucial insights into its *in vivo* organisation and function in acid stress response.

A recent systematic analysis of polymerisation-disrupting mutations in yeast [14] demonstrated that altering the surface chemistry of a dihedral symmetric protein can strongly influence its supramolecular self-assembly and intracellular localisation. Importantly, our work quantitatively confirms the preferential peripheral distribution of WT LdcI clusters, inferred from visual inspection in our earlier studies [17]. Moreover, even mutant LdcI clusters, though significantly smaller, exhibit a tendency for peripheral localisation. This suggests that, despite diminished filamentation efficiency, the mutant LdcI retains the ability to localise to specific cellular sites, likely reflecting a cellular strategy to concentrate LdcI where it is most needed. Such assembly would enhance enzymatic activity thereby tuning the substrate-product turnover at these sites. The exact nature of these peripheral sites and their specific role in acid stress response remain intriguing open questions.

The enzymatic reaction of LdcI converts a proton, lysine and water molecules into cadaverine and carbon dioxide. Thus, it might be advantageous for the *E. coli* cell to position LdcI near the inner membrane lysine permease LysP and/or the lysine/cadaverine aniporter CadB. While to our knowledge the localisation of LysP and CadB has not yet been investigated by SMLM, wide field fluorescence images of *E. coli* MG1655 strains harboring chromosomally encoded mCherry-LysP or CadB-eGFP fusions did not show any indication of a patchy distribution [33]. Alternatively, considering that LdcI directly consumes protons, it might localise to regions of high local proton concentration, such as cardiolipin-enriched lipid microdomains, known to be unevenly distributed in the inner membrane as mobile patches [42–44]. These microdomains are involved in localisation, assembly, and regulation of respiratory complexes, which couple electron transport with proton export, generating the proton motive force (PMF) required for cellular energy production [43, 45–47]. Under mild acid stress, *E. coli* upregulates components of its electron transport chain to counteract intracellular acidification by directly increasing proton export [2, 48, 49]. Interestingly, partially assembled respiratory complex I was reported to co-purify with LdcI [50]. In addition, complex I clusters within the *E. coli* inner membrane are reminiscent of the LdcI clusters observed here, albeit those experiments were conducted under aerobic, non-acidic conditions [51]. Thus, one could imagine that LdcI may polymerise near complex I, benefiting from elevated proton concentrations to counteract acid stress by consuming protons intracellularly while complex I exports them. Furthermore, a cardiolipin-binding protein ViaA and its partner RavA, which strongly interacts with LdcI, were described to associate with *E. coli* complex I and fumarate reductase, and to localise to the inner membrane, although the localisation has so far only been investigated under overexpression conditions [39, 52–55]. It is worth noting, however, that our current experiments did not provide clear evidence of preferential localisation of the LdcI clusters to the cell poles or division plane, as expected for cardiolipin-enriched patches that should favor regions with negative membrane curvature [56]. Thus, while we have now clearly established that LdcI clustering within *E. coli* cells is driven by its polymerisation, essential to mount an efficient acid stress response, the next step towards fully understanding LdcI function would be to directly investigate its colocalisation with respiratory complexes, cardiolipin, RavA and ViaA, LysP, CadB, and other potentially relevant systems that may contribute to the positioning of LdcI clusters at these specific sites at the cell periphery.

For a neutralophilic bacterium, *E. coli* exhibits a remarkably robust ability to withstand a wide range of acid stress conditions, a testament to the extensive repertoire of acid resistance pathways within the cell and the efficient division of labor among them. Our study corroborates recent findings on the differential expression of the Cad system across a population of acid-stressed *E. coli* [33, 57], underscores the value of quantitative SMLM imaging in unraveling the sophisticated strategies that enable bacterial survival under challenging environmental conditions, and prompts further exploration of enzyme filamentation as a key element in stress responses.

## Acknowledgments

We thank staff of the IBS M4D platform, and in particular Oleksandr Glushonkov, for their support and advice. We are grateful to Clarissa Liesche for her initial work on the LdcI clustering, which motivated the present study, and to Alain Roussel and Aline Desmyter for the anti-LdcI nanobody production. This work was funded by the European Union’s Horizon 2020 research and innovation programme under grant agreement 647784, and the CBH-EUR graduate school (ANR-17-EURE-0003) and the IDEX IRGA 2023 of the University Grenoble Alpes. We acknowledge the IBS integration into the Interdisciplinary Research Institute of Grenoble (IRIG, CEA). The optical imaging was carried out on the M4D imaging platform of the Grenoble Instruct-ERIC center (ISBG; UAR 3518 CNRS-CEA-UGA-EMBL) within the Grenoble Partnership for Structural Biology (PSB), supported by FRISBI (ANR-10-INBS-0005-02) and GRAL, financed within the University Grenoble Alpes graduate school (Ecoles Universitaires de Recherche) CBH-EUR-GS (ANR-17-EURE-0003). The nanobody generation platform of the AFMB laboratory (Marseille, France) was supported by the French Infrastructure for Integrated Structural Biology FRISBI) ANR-10-INSB-05-01. M.K. is supported by a CEA CFR PhD fellowship.

## Data availabilty

A tabular file including all statistics on cluster volumes and localisation, as well as liquid culture pH and OD values has been uploaded to a Zenodo repository (https://doi.org/10.5281/zenodo.15094803). The repository includes an all-in-one python script for cluster volume and location determination as described in this manuscript to be used on output FOCAL3D .txt files and a .yml file for setup of a related conda environment to run the program.

## Notes

### Competing Interest Statement

The authors have declared no competing interest.

## References

1. Fang FC, Frawley ER, Tapscott T, Vázquez-Torres A. Bacterial stress responses during host infection. Cell host & microbe. 2016;20(2):133–143.

2. Kanjee U, Houry WA. Mechanisms of acid resistance in Escherichia coli. Annual review of microbiology. 2013;67(1):65–81.

3. Foster JW. Escherichia coli acid resistance: tales of an amateur acidophile. Nature Reviews Microbiology. 2004;2(11):898–907.

4. Michael AJ. Polyamine function in archaea and bacteria. Journal of Biological Chemistry. 2018;293(48):18693–18701.

5. De Biase D, Lund PA. The Escherichia coli acid stress response and its significance for pathogenesis. Advances in applied microbiology. 2015;92:49–88.

6. Brameyer S, Schumacher K, Kuppermann S, Jung K. Division of labor and collective functionality in Escherichia coli under acid stress. Communications Biology. 2022;5(1):327.

7. Watson N, Dunyak D, Rosey E, Slonczewski J, Olson E. Identification of elements involved in transcriptional regulation of the Escherichia coli cad operon by external pH. Journal of bacteriology. 1992;174(2):530–540.

8. Rauschmeier M, Schüppel V, Tetsch L, Jung K. New insights into the interplay between the lysine transporter LysP and the pH sensor CadC in Escherichia coli. Journal of molecular biology. 2014;426(1):215–229.

9. Küper C, Jung K. CadC-mediated activation of the cadBA promoter in Escherichia coli. Journal of molecular microbiology and biotechnology. 2005;10(1):26–39.

10. Brameyer S, Rösch TC, El Andari J, Hoyer E, Schwarz J, Graumann PL, et al. DNA-binding directs the localization of a membrane-integrated receptor of the ToxR family. Communications biology. 2019;2(1):4.

11. Moreau PL. The lysine decarboxylase CadA protects Escherichia coli starved of phosphate against fermentation acids. Journal of bacteriology. 2007;189(6):2249–2261.

12. Kanjee U, Gutsche I, Alexopoulos E, Zhao B, El Bakkouri M, Thibault G, et al. Linkage between the bacterial acid stress and stringent responses: the structure of the inducible lysine decarboxylase. The EMBO journal. 2011;30(5):931–944.

13. Ma W, Chen K, Li Y, Hao N, Wang X, Ouyang P. Advances in cadaverine bacterial production and its applications. Engineering. 2017;3(3):308–317.

14. Garcia Seisdedos H, Levin T, Shapira G, Freud S, Levy ED. Mutant libraries reveal negative design shielding proteins from supramolecular self-assembly and relocalization in cells. Proceedings of the National Academy of Sciences. 2022;119(5):e2101117119.

15. Park CK, Horton NC. Structures, functions, and mechanisms of filament forming enzymes: a renaissance of enzyme filamentation. Biophysical reviews. 2019;11(6):927–994.

16. Garcia-Seisdedos H, Empereur-Mot C, Elad N, Levy ED. Proteins evolve on the edge of supramolecular self-assembly. Nature. 2017;548(7666):244–247.

17. Jessop M, Liesche C, Felix J, Desfosses A, Baulard M, Adam V, et al. Supramolecular assembly of the Escherichia coli LdcI upon acid stress. Proceedings of the National Academy of Sciences. 2021;118(2):e2014383118. doi:10.1073/pnas.2014383118.

18. Jessop M, Huard K, Desfosses A, Tetreau G, Carriel D, Bacia-Verloop M, et al. Structural and biochemical characterisation of the Providencia stuartii arginine decarboxylase shows distinct polymerisation and regulation. Communications Biology. 2022;5(1):317.

19. Carsten A, Wolters M, Aepfelbacher M. Super-resolution fluorescence microscopy for investigating bacterial cell biology. Molecular Microbiology. 2024;121(4):646–658.

20. Pierson JA, Yang JE, Wright ER. Recent advances in correlative cryo-light and electron microscopy. Current Opinion in Structural Biology. 2024;89:102934.

21. Prouteau M, Loewith R. Regulation of cellular metabolism through phase separation of enzymes. Biomolecules. 2018;8(4):160.

22. Datsenko KA, Wanner BL. One-step inactivation of chromosomal genes in Escherichia coli K-12 using PCR products. Proceedings of the National Academy of Sciences. 2000;97(12):6640–6645.

23. Trouve J, Glushonkov O, Morlot C. Metabolic biorthogonal labeling and dSTORM imaging of peptidoglycan synthesis in Streptococcus pneumoniae. STAR protocols. 2021;2(4):101006.

24. Rust MJ, Bates M, Zhuang X. Sub-diffraction-limit imaging by stochastic optical reconstruction microscopy (STORM). Nature methods. 2006;3(10):793–796.

25. Heilemann M, Van De Linde S, Schüttpelz M, Kasper R, Seefeldt B, Mukherjee A, et al. Subdiffraction-resolution fluorescence imaging with conventional fluorescent probes. Angewandte Chemie-International Edition. 2008;47(33).

26. Tokunaga M, Imamoto N, Sakata-Sogawa K. Highly inclined thin illumination enables clear single-molecule imaging in cells. Nature methods. 2008;5(2):159–161.

27. Huang B, Wang W, Bates M, Zhuang X. Three-dimensional super-resolution imaging by stochastic optical reconstruction microscopy. Science. 2008;319(5864):810–813.

28. Wang Y, Schnitzbauer J, Hu Z, Li X, Cheng Y, Huang ZL, et al. Localization events-based sample drift correction for localization microscopy with redundant cross-correlation algorithm. Optics express. 2014;22(13):15982–15991.

29. Ester M, Kriegel HP, Sander J, Xu X, et al. A density-based algorithm for discovering clusters in large spatial databases with noise. 1996;96(34):226–231.

30. Ripley BD. The second-order analysis of stationary point processes. Journal of applied probability. 1976;13(2):255–266.

31. Doose S. LOCAN: a python library for analyzing single-molecule localization microscopy data. Bioinformatics. 2022;38(9):2670–2672.

32. Nino DF, Djayakarsana D, Milstein JN. FOCAL3D: A 3-dimensional clustering package for single-molecule localization microscopy. PLoS computational biology. 2020;16(12):e1008479.

33. Brameyer S, Hoyer E, Bibinger S, Burdack K, Lassak J, Jung K. Molecular design of a signaling system influences noise in protein abundance under acid stress in different gammaproteobacteria. Journal of Bacteriology. 2020;202(16):10–1128.

34. Andronov L, Orlov I, Lutz Y, Vonesch JL, Klaholz BP. ClusterViSu, a method for clustering of protein complexes by Voronoi tessellation in super-resolution microscopy. Scientific reports. 2016;6(1):24084.

35. Levet F, Hosy E, Kechkar A, Butler C, Beghin A, Choquet D, et al. SR-Tesseler: a method to segment and quantify localization-based super-resolution microscopy data. Nature methods. 2015;12(11):1065–1071.

36. Hyun Y, Kim D. Recent development of computational cluster analysis methods for single-molecule localization microscopy images. Computational and Structural Biotechnology Journal. 2023;21:879–888.

37. Kandiah E, Carriel D, Garcia PS, Felix J, Banzhaf M, Kritikos G, et al. Structure, function, and evolution of the Pseudomonas aeruginosa lysine decarboxylase LdcA. Structure. 2019;27(12):1842–1854.

38. Felix J, Siebert C, Ducassou JN, Nigou J, Garcia PS, Fraudeau A, et al. Structural and functional analysis of the Francisella lysine decarboxylase as a key actor in oxidative stress resistance. Scientific Reports. 2021;11(1):972.

39. Jessop M, Arragain B, Miras R, Fraudeau A, Huard K, Bacia-Verloop M, et al. Structural insights into ATP hydrolysis by the MoxR ATPase RavA and the LdcI-RavA cage-like complex. Communications Biology. 2020;3(1):46.

40. Garcia-Seisdedos H, Villegas JA, Levy ED. Infinite assembly of folded proteins in evolution, disease, and engineering. Angewandte Chemie International Edition. 2019;58(17):5514–5531.

41. Hvorecny KL, Kollman JM. Greater than the sum of parts: mechanisms of metabolic regulation by enzyme filaments. Current opinion in structural biology. 2023;79:102530.

42. Haines TH, Dencher NA. Cardiolipin: a proton trap for oxidative phosphorylation. FEBS letters. 2002;528(1-3):35–39.

43. Arias-Cartin R, Grimaldi S, Arnoux P, Guigliarelli B, Magalon A. Cardiolipin binding in bacterial respiratory complexes: structural and functional implications. Biochimica et Biophysica Acta (BBA)-Bioenergetics. 2012;1817(10):1937–1949.

44. Magalon A, Alberge F. Distribution and dynamics of OXPHOS complexes in the bacterial cytoplasmic membrane. Biochimica et Biophysica Acta (BBA)-Bioenergetics. 2016;1857(3):198–213.

45. Pfeiffer K, Gohil V, Stuart RA, Hunte C, Brandt U, Greenberg ML, et al. Cardiolipin stabilizes respiratory chain supercomplexes. Journal of biological chemistry. 2003;278(52):52873–52880.

46. Jussupow A, Di Luca A, Kaila VR. How cardiolipin modulates the dynamics of respiratory complex I. Science advances. 2019;5(3):eaav1850.

47. Fry M, Green DE. Cardiolipin requirement for electron transfer in complex I and III of the mitochondrial respiratory chain. Journal of Biological Chemistry. 1981;256(4):1874–1880.

48. Maurer LM, Yohannes E, Bondurant SS, Radmacher M, Slonczewski JL. pH regulates genes for flagellar motility, catabolism, and oxidative stress in Escherichia coli K-12. Journal of bacteriology. 2005;187(1):304–319.

49. Slonczewski JL, Fujisawa M, Dopson M, Krulwich TA. Cytoplasmic pH measurement and homeostasis in bacteria and archaea. Advances in microbial physiology. 2009;55:1–317.

50. Erhardt H, Steimle S, Muders V, Pohl T, Walter J, Friedrich T. Disruption of individual nuo-genes leads to the formation of partially assembled NADH: ubiquinone oxidoreductase (complex I) in Escherichia coli. Biochimica et Biophysica Acta (BBA)-Bioenergetics. 2012;1817(6):863–871.

51. Llorente-Garcia I, Lenn T, Erhardt H, Harriman OL, Liu LN, Robson A, et al. Single-molecule in vivo imaging of bacterial respiratory complexes indicates delocalized oxidative phosphorylation. Biochimica et Biophysica Acta (BBA)-Bioenergetics. 2014;1837(6):811–824.

52. Snider J, Gutsche I, Lin M, Baby S, Cox B, Butland G, et al. Formation of a distinctive complex between the inducible bacterial lysine decarboxylase and a novel AAA+ ATPase. Journal of Biological Chemistry. 2006;281(3):1532–1546.

53. Wong KS, Snider JD, Graham C, Greenblatt JF, Emili A, Babu M, et al. The MoxR ATPase RavA and its cofactor ViaA interact with the NADH: ubiquinone oxidoreductase I in Escherichia coli. PloS one. 2014;9(1):e85529.

54. Wong KS, Bhandari V, Janga SC, Houry WA. The RavA-ViaA chaperone-like system interacts with and modulates the activity of the fumarate reductase respiratory complex. Journal of Molecular Biology. 2017;429(2):324–344.

55. Felix J, Bumba L, Liesche C, Fraudeau A, Rébeillé F, El Khoury JY, et al. The AAA+ ATPase RavA and its binding partner ViaA modulate E. coli aminoglycoside sensitivity through interaction with the inner membrane. Nature communications. 2022;13(1):5502.

56. Renner LD, Weibel DB. Cardiolipin microdomains localize to negatively curved regions of Escherichia coli membranes. Proceedings of the National Academy of Sciences. 2011;108(15):6264–6269.

57. Schumacher K, Brameyer S, Jung K. Bacterial acid stress response: From cellular changes to antibiotic tolerance and phenotypic heterogeneity. Current Opinion in Microbiology. 2023;75:102367.

