## Supplementary information for "Filamentation-driven peripheral clustering of the inducible lysine decarboxylase is crucial for *E. coli* acid stress response"

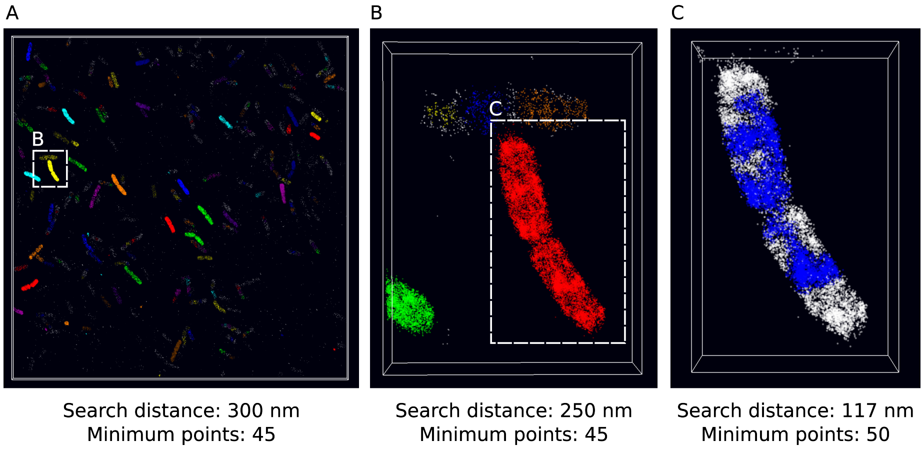


**S1 Fig. Segmentation of single cells from the full recorded volume using DBSCAN clustering.** One colour represents a single cluster as determined by DBSCAN. A: Full 3D image segmented with DBSCAN clustering. B: A few cells from A which were segmented into the same cluster by DBSCAN on the global scale but are clearly different cells. In a second step the example subvolume C, outlined with dashed lines, was extracted and subjected to clustering again with slightly adjusted parameters to assure better cell separation. C: A pair of recently divided cells which could not accurately be separated by DBSCAN because the largest cluster (blue) detected before the entire volume of the cells was segmented, spans both cells. These cells were used as a 'two-cell file' for cluster volume determination but excluded from determination of distance ratios (see Methods), polar location determination (S2 Fig.) and total localisation number determination per cell (S4 Fig.).

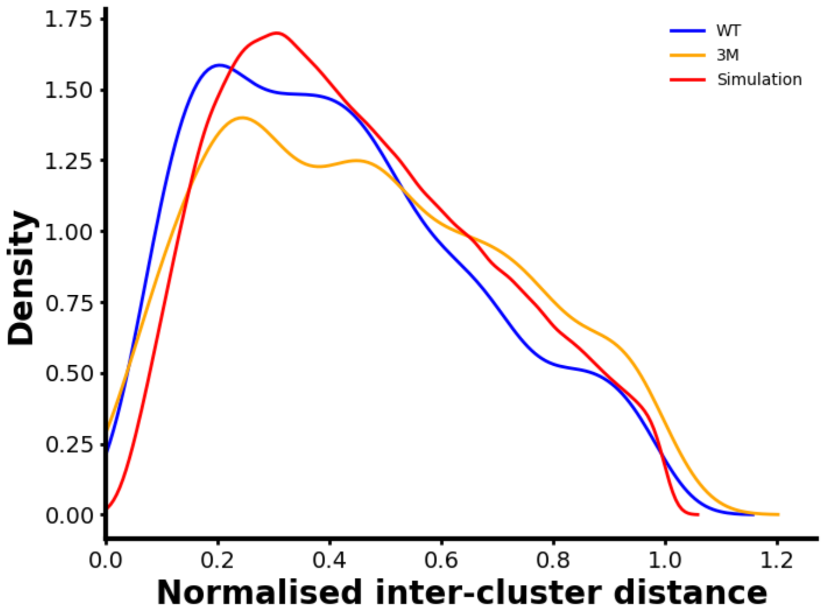


**S2 Fig. Kernel density estimation of the distribution of inter-cluster distances in *E. coli* WT and 3M cells.** All inter-cluster distances were determined between cluster centroids. Normalisation to the longest distance measured in each cell. The longest distance of each cell (always 1) was omitted from this representation. The simulation describes the distances measured for 100 randomly placed test-centroids per cell in the cell volume.

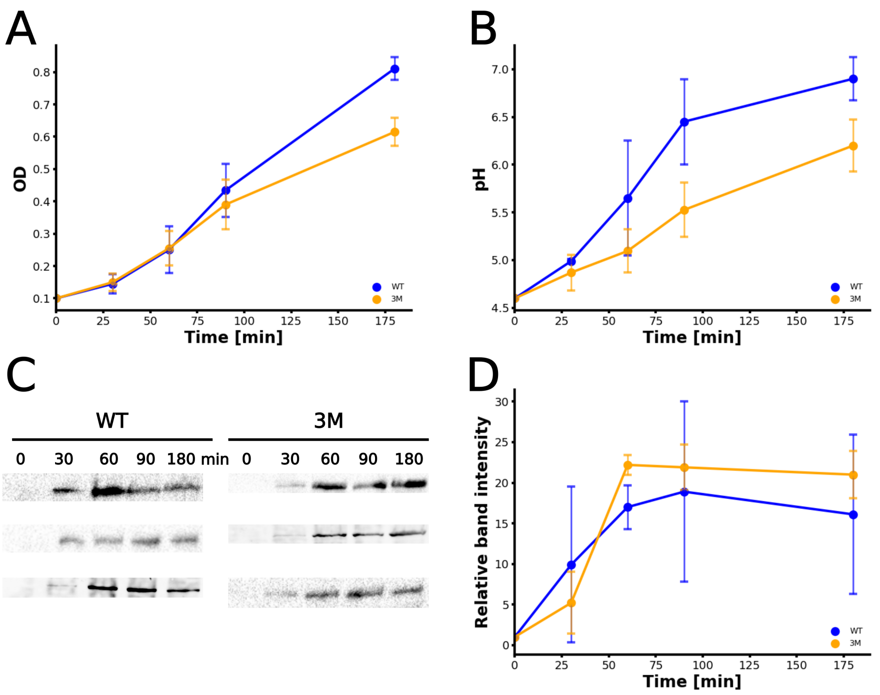


**S3 Fig. Impact of LdcI triple mutant on *E. coli* acid stress response.** A: Growth curves of *E. coli* WT and 3M liquid culture under acid stress (inital pH 4.6). B: pH of the external medium during *E. coli* WT or 3M growth in a pH shift experiment. C: Triplicate of western blot with full cell extract from WT and 3M cultures over the first 180 min after the pH drop to 4.6. D: Average band intensities retrieved from the blots shown in C. Values were normalised to intensities measured for the LdcI band at 0 min on each blot individually.

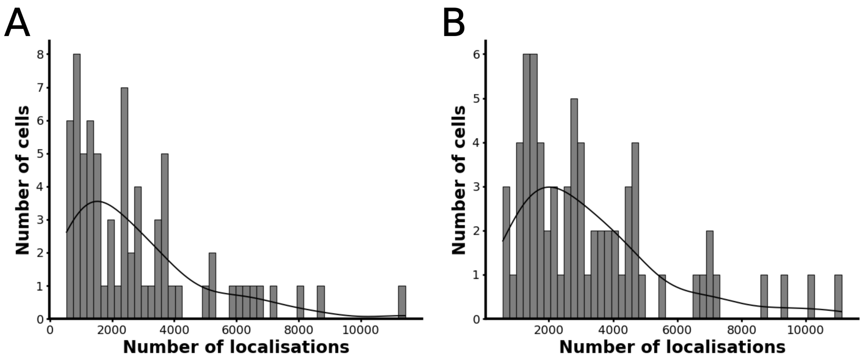


**S4 Fig. Distribution of the total number of cells per number of recorded intra-cell localisations.** A: WT cells. B: 3M cells.

**S1 File. Supplementary tables.** Page 1: Statistics on cluster volumes and localisation. Tabular file containing all cluster volumes and distances ratios (everywhere applicable, see Methods), as well as raw data on liquid culture OD and pH values. Page 2: Table listing plasmids, strains and primers used for creating the 3M ldcI chromosomal mutant.
